# A Novel Non-Invasive Epithelial Ovarian Cancer Mouse Model Of Hyperthermic Intraperitoneal Chemotherapy (HIPEC)

**DOI:** 10.1101/2021.05.16.444343

**Authors:** Zahraa Alali, Max P. Horowitz, Danielle Chau, Lexie Trestan, Jing Hao, Peng Qi, Emily L. Esakov, Robert L. DeBernardo, Jennifer S. Yu, Ofer Reizes

## Abstract

**Background:** Hyperthermic intraperitoneal chemotherapy (HIPEC) in combination with interval cytoreductive surgery increases the overall survival of epithelial ovarian cancer (EOC) patients with advanced disease. Despite its proven benefits, the mechanism by which HIPEC extends overall survival remains unknown and current strategies to optimize HIPEC are therefore limited. A major challenge is the lack of a robust and streamlined model to investigate the mechanisms underlying HIPEC efficacy.

**Objective:** To introduce a novel murine model that can be used to enhance our understanding of HIPEC therapy.

**Method:** ID8-luc, an EOC mouse cell line, is inoculated into immunocompetent C57BL/6J mice intraperitoneally. Once tumor is detected by In Vivo Imaging System (IVIS), cisplatin (5 mg/kg) is injected intraperitoneally and superficial hyperthermia of 40°C is applied to the animal’s abdomen and pelvis using an FDA-approved hyperthermia unit (BSD500) for 20 minutes. To validate the model, four treatment conditions were tested: cisplatin and hyperthermia, cisplatin and normothermia, vehicle and hyperthermia, and vehicle and normothermia. Tumor growth was assessed over the course of treatment using IVIS optical spectrum.

**Results:** Tumor growth in mice treated with hyperthermic cisplatin was significantly suppressed compared to mice treated with normothermic cisplatin (p < 0.05). No significant differences in tumor growth were observed in the hyperthermic vehicle and normothermic vehicle groups.

**Conclusions:** We developed an innovative noninvasive mouse model of HIPEC. Similar to patients with advanced ovarian cancer who are treated with HIPEC at the time of interval cytoreductive surgery, our model demonstrates that hyperthermia enhances the inhibitory effect of cisplatin on intraperitoneal tumor growth. Development of this murine model provides an opportunity to elucidate the mechanisms underlying HIPEC and offer an opportunity to test adjunct treatments in a pre-clinical setting to enhance the utility of HIPEC.

## INTRODUCTION

Advanced epithelial ovarian cancer (EOC) carries a poor prognosis despite standard of care therapy involving the combination of surgery and systemic chemotherapy. The majority of patients present with advanced stage disease that has disseminated throughout the abdominal cavity, often necessitating systemic chemotherapy upfront to reduce the morbidity of aggressive surgery. Typical 5-year overall survival of for these patients with advanced disease is approximately 30% [1]. Ovarian cancer remains a leading cause of cancer death in women and unfortunately, there have been few improvements in oncologic outcomes for patients with advanced ovarian cancer over the past 15 years [2, 3].

HIPEC therapy has been utilized clinically with success for several non-gynecologic malignancies [5-11]. In 2018, van Driel et al. published a randomized controlled trial of patients with advanced ovarian cancer who had undergone optimal interval debulking surgery following neoadjuvant platinum/taxane-based chemotherapy [4]. Patients were randomized to either surgery alone followed by adjuvant chemotherapy or surgery with concurrent hyperthermic intraperitoneal chemotherapy (HIPEC) followed by adjuvant chemotherapy. The outcome of the study indicates that the addition of HIPEC at the time of interval debulking surgery conferred a statistically significant benefit in both recurrence-free survival (3.5-month benefit) as well as overall survival (11.9-month benefit) [4].

HIPEC therapy has similarly been utilized clinically for several intraperitoneal malignancies [5-11]. Despite this success in extending survival for patients with advanced ovarian cancer, the mechanism by which HIPEC improves patient outcomes remains unknown. This poses a significant barrier to optimizing HIPEC protocols for patient benefit. There exists a need for a reliable animal model to interrogate this mechanism further. Currently available HIPEC models are cumbersome with complex protocols for inflow/outflow, lack reproducibility, and are typically done in animal models in which genetics are not readily manipulable [12-17]. The large majority of these models utilize GI cancer cell lines and do not confirm the presence of peritoneal carcinomatosis prior to the initiation of HIPEC [12-17]. Furthermore, few models exist that allow for safe and efficient delivery of heated chemotherapy in multiple mice per procedure.

We sought to develop an animal model of HIPEC that could overcome these limitations, while allowing for further interrogation of the mechanisms that mediate the beneficial effects of HIPEC and in vivo studies of combinations of therapeutics with HIPEC. Since cisplatin is used clinically in the treatment of EOC and has also been used successfully in animal models of hyperthermia [21], we decided to use cisplatin in our non-invasive mouse model of HIPEC. We show here the development of this novel approach for the study of HIPEC in a murine model of ovarian cancer. Using *in vivo* imaging (IVIS), we show that the addition of hyperthermia delivered via heated probes and intraperitoneal injection of cisplatin leads to significantly decreased tumor growth compared to normothermic cisplatin. In conclusion, this is the first non-invasive animal model of HIPEC for advanced ovarian cancer in a platform amenable to genetic and pharmacologic manipulation.

## MATERIALS AND METHODS

The experiments were conducted in accordance with the National Institute of Health (NIH) Guide for the Care and Use of Laboratory Animals. All protocols were approved by Cleveland Clinic IACUC.

### Cell culture

Mouse epithelial ovarian cancer expressing luciferase (ID8-luc) were generated in the lab as previously described [22, 23]. ID8-luc were cultured in in Dulbecco Modified 345 Eagle Medium (DMEM) media containing heat inactivated 5% FBS (Atlas Biologicals Cat # F-0500-D, 346 Lot F31E18D1) and grown under standard conditions. At confluence, cells were washed, trypsinized, and centrifuged to form a cell pellet. Cell pellets were resuspended in FBS-free DMEM media (Dulbecco Modified 345 Eagle Medium), and this preparation was used for injection.

### Epithelial Ovarian Cancer EOC induction in mice

6-8 week old female C57BL/6J mice from Jackson Laboratories (Bar Harbor, ME) were injected intraperitoneally with 5×10^6^ ID8-luc cells (in 0.3 ml FBSfree DMEM media). The mice were housed in standard housing in accordance with Cleveland Clinic Lerner Research Institute Biological Resources Unit protocols.

### In vivo imaging

Two weeks after cell inoculation, mice were injected with D-luciferin intraperitoneally (Goldbio LUCK-1G, 150mg/kg in 200mL) and anesthetized with inhaled isoflurane. After 10 minutes, mice were placed in IVIS Lumina (PerkinElmer) system to obtain bioluminescence images. The analysis of the images was done using Living Image Software, and the intensity of the signal (total flux) was recorded. This process was repeated weekly until endpoint.

### HIPEC application

To apply heat to the abdomen and pelvis, mice were placed in prone position with their abdomen and pelvis above a water circulation system with enclosed applicator bolus with water temperature set at 40°C. Hyperthermia was delivered using the FDA-approved BSD500 instrument (Pyrexar). The peritoneal cavity was heated using a single microwave antenna (915 MHz) affixed under the abdominopelvic region of each mouse (Figure 1). Non-invasive wire thermistors that control the microwave power to monitor real-time temperature were placed adjacent to the microwave antennae for each mouse. Mice were monitored for signs of hyperventilation or stress.

**Figure 1.**
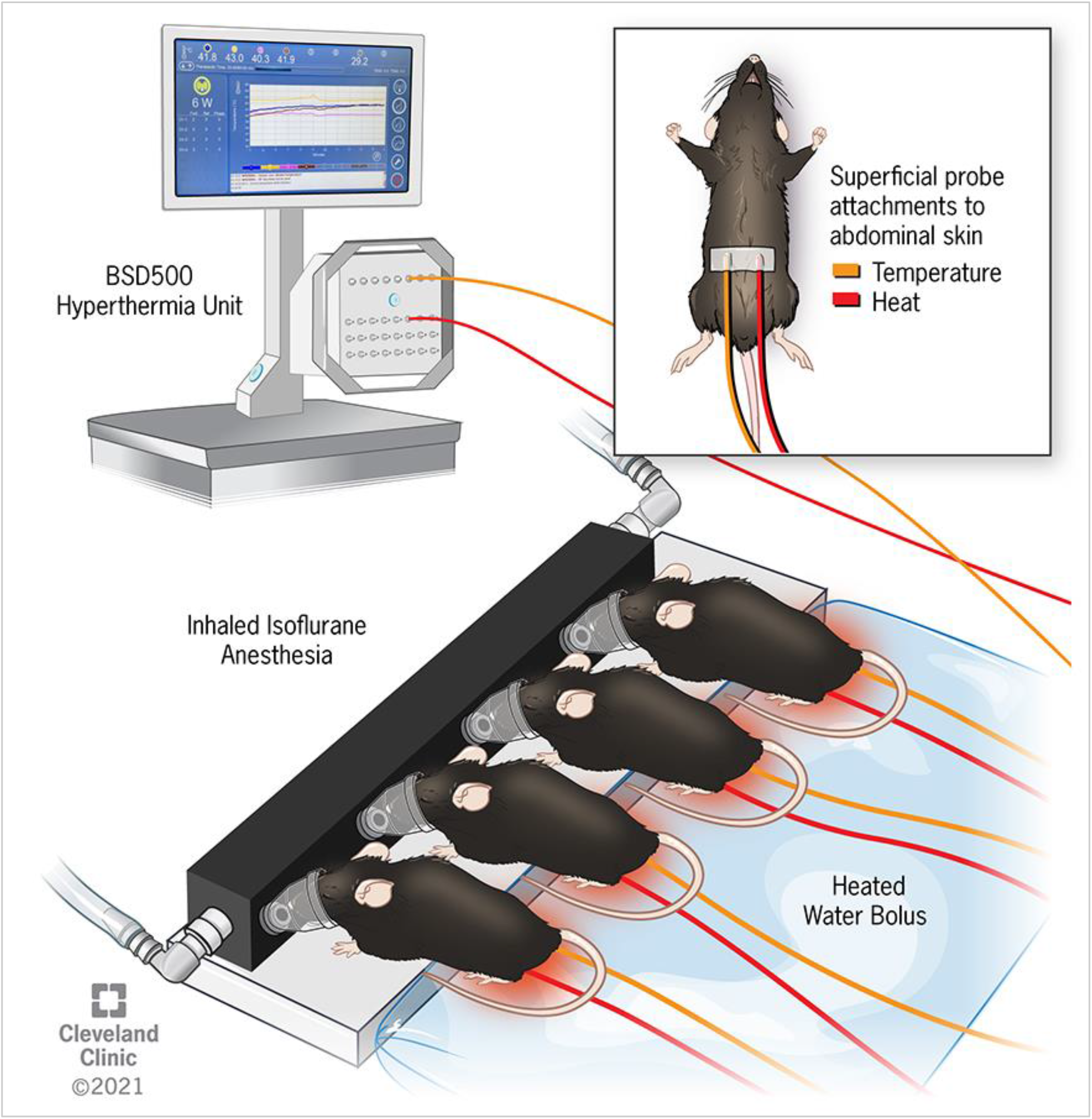
Schematic representation of device for induction of Hyperthermic Intraperitoneal Chemotherapy (HIPEC) in mice. a microwave antenna and thermistor wire are superficially attached to the abdomen. Mice are anesthetized with inhaled isoflurane and placed in a prone position on a water bolus. Heat at 40-42°C is administered for 90-20 minutes.

### Statistical analysis

The total photon flux of each time point, calculated by IVIS, was subtracted from the pre-HIPEC measurement to evaluate the change. The data were analyzed with Prism (GraphPad Prism 8) using one-way ANOVA analysis; p<0.05 was considered as statistically significant.

## RESULTS

### Optimization of Hyperthermia Treatment in Immune Competent C57BL/6J Mice

To evaluate the safety and the optimal time and temperature of the procedure (Figure 1), tumor-free C57BL/6J mice were subjected to several conditions in three separate trials: [1] 42°C for 90 minutes, [2] 42°C for 40 minutes, [3] 40°C for 20 minutes.

The first trial was discontinued after 50 minutes due to a mouse mortality and signs of respiratory distress (e.g. hyperventilation) in the remaining mice. In the second trial, all mice showed signs of respiratory distress after 30 minutes and the study was discontinued. Adjustments were made in the third trial to account for these findings; the temperature was reduced to 40°C and continued for 20 minutes. These changes resulted in safe induction of heat without complication or mortality. Therefore, we used these conditions with tumor-bearing mice.

### Impact of Cisplatin Treatment in the Presence and Absence of Hyperthermia in Murine EOC

C57Bl/6J mice were injected intraperitoneally with ID8-Luc cells, a validated mouse murine EOC model with similarity to high grade serous ovarian cancer. A schematic representation of the treatment protocol is shown in Figure 2A. After two weeks, IVIS imaging demonstrated that mice developed abdominoperitoneal metastases. The observed metastases are similar to those observed in patients with advanced ovarian cancer [24]. Mice were randomized to receive an IP injection of either 200 ul of saline (vehicle) or cisplatin (5 mg/kg). Mice were subsequently anesthetized and immobilized on the hyperthermic device. The abdominopelvic cavities were heated to either 40°C or not heated (37°C) for a period of 20 minutes while under anesthesia. An underlying water circulation system set at 40°C was employed for even distribution of heat therapy and to ensure stabilization of body temperature. Superficial temperatures of the animals were measured throughout the 20 minutes and were recorded and stable body temperature was noted for the duration of hyperthermic treatment (Figure 1).

**Figure 2.**
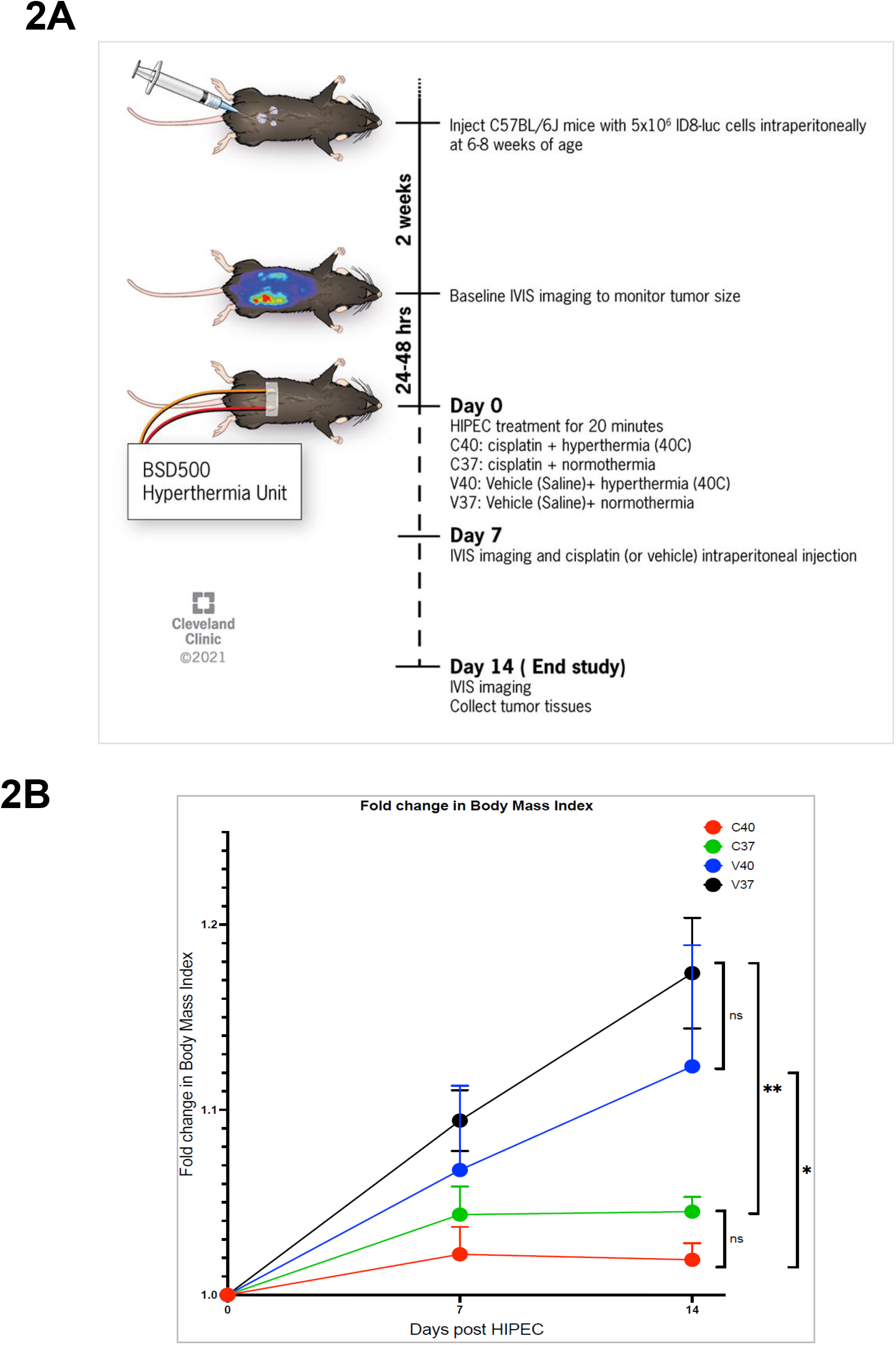
Treatment of EOC mouse model with Cisplatin in Presence and Absence of Hyperthermia. **(A)** HIPEC protocol for mice bearing peritoneal Epithelial Ovarian Cancer EOC. **(B)** The weight of mice were measured on the day of HIPEC (Pre-HIPEC, Day 0), and then weekly after HIPEC (Day 7, and Day 14 post HIPEC). Graphed data, means ± SEM; *, *P* < 0.05; **, *P* < 0.01; ***, *P* < 0.001. C40: mice treated cisplatin (5mg/kg) and hyperthermia at 40°C (n=4); C37: mice treated with cisplatin (5mg/kg) and normothermia (n=5). V37 Mice treated with Saline and normothermia (n=5). V40: mice treated with saline with hyperthermia (n=4).

There were no animal deaths during the procedure and mice tolerated the heat with no observeable side effects. Mice were observed daily per the Biological Resources Unit protocol for the duration of the study and weighed on a weekly basis. There was no significant dehydration or abdominal wounds noted across all treatment groups. However, mice treated with normothermic (C37) or hyperthermic cisplatin (C40) exhibited significant attenuation of body mass (*P* < 0.05) compared to normothermic (V37) or hyperthermic (V40) saline treated mice suggestive of reduced tumor burden (Figure 2B).

Cisplatin or saline was administered on a weekly basis and mice were imaged using IVIS to track tumor growth. Change in total flux was calculated for each treatment group. Tumor growth based on change in bioluminescence signal intensity, was significantly reduced (P< 0.05) in mice treated with cisplatin (5mg/kg) and hyperthemia at 40°C (C40) group, at day 7 and day 14 post HIPEC compared to mice treated with cisplatin (5mg/kg) and normothermia (C37) group (Figure 3A, 3B). Vehicle-treated mice exposed to hyperthermia exhibited no significant difference in tumor growth compared to normothermic mice (Figure 3A,3B).

**Figure 3.**
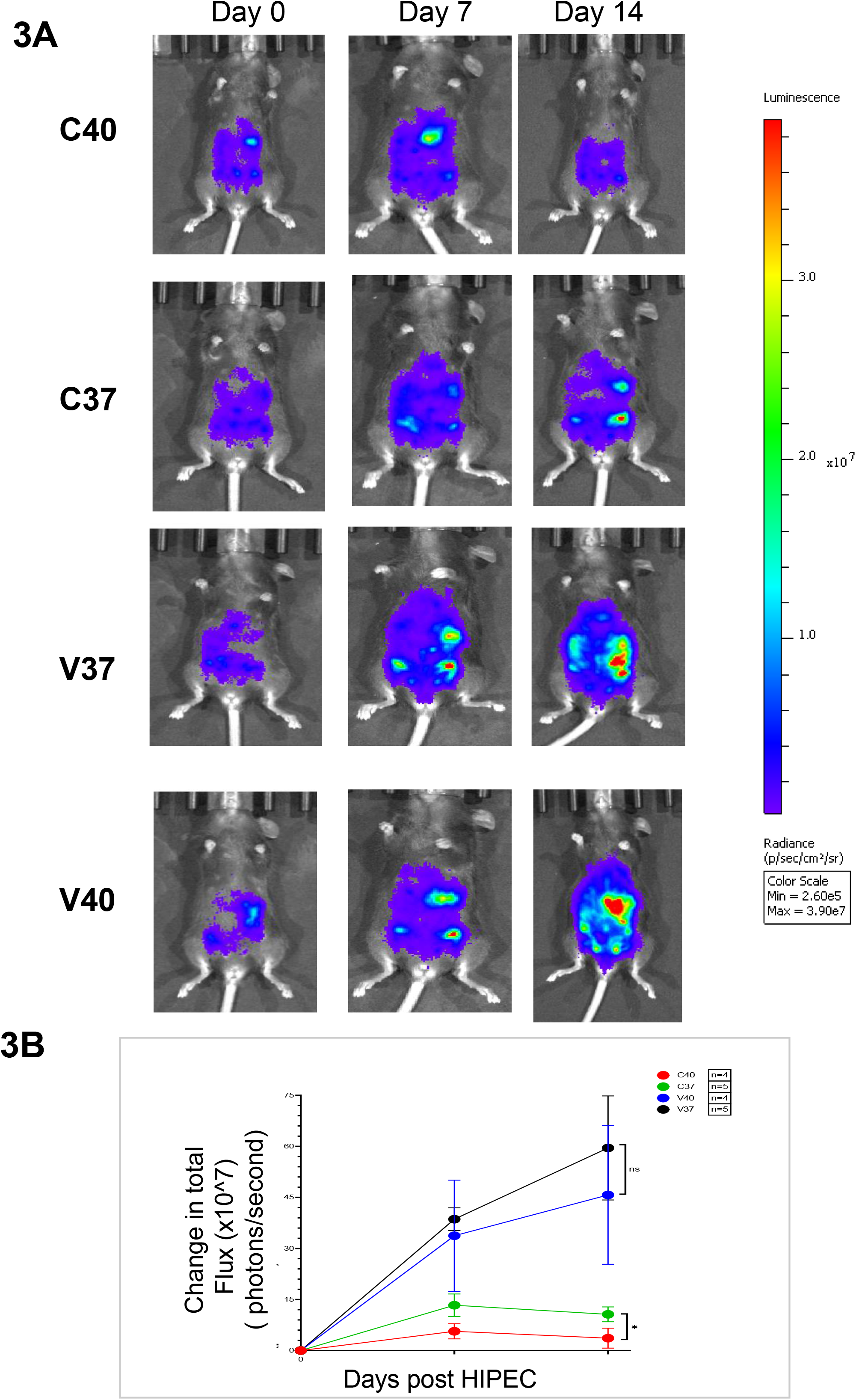
In vivo imaging of C57BL/6J mice injected with ID8-Luc cells using IVIS bioluminescence imaging system. (A) Representative images of mice from baseline (day 0) to 2 weeks post-HIPEC. Increase in intensity of bioluminescence signal represents growth of ovarian tumors in C57BL/6J mice. **(B)** Change in bioluminescence signal (total flux) at day 0 (pre-HIPEC) and at day 7 and day 14 post HIPEC. Graphed data, means ± SEM; *, *P* < 0.05.(B) Change in bioluminescence signal (total flux) at day 0 (pre-HIPEC) and at day 7 and day 14 post HIPEC. Day 0.Graphed data, means ± SEM; *, *P* < 0.05. C40: mice treated cisplatin (5mg/kg) and hyperthermia at 40°C (n=4); C37: mice treated with cisplatin (5mg/kg) and normothermia (n=5). V37 Mice treated with Saline and normothermia (n=5). V40: mice treated with saline with hyperthermia (n=4).

## DISCUSSION

As the proven clinical benefit of HIPEC drives ongoing clinical trials for advanced EOC patients [25], the mechanism of action remains unknown. This limits the ability to develop combinatorial therapeutics to further enhance the clinical benefit of HIPEC. As such, there is a need to develop a pre-clinical model by which to interrogate the mechanism of HIPEC in a safe and efficient manner. Previous animal trials have utilized an invasive approach in inducing HIPEC, limiting the ability to perform mechanistic or pre-clinical evaluation of multiple therapeutic strategies. For instance, G. Miailhe et al. used an open abdomen procedure that showed less stable temperature throughout treatment and caused bleeding. They then moved to a less invasive procedure with closed abdomen and a circuit system with inflow and outflow catheter to circulate the heated oxaliplatin at 53°C, but maintain an inside abdomen temperture at 43°C [12]. Such models are fraught with technical challenges that undermine their utility for wider pre-clinical use.

Our model, by shifting the paradigm from an invasive to a non-invasive approach, represents a technological advance in interrogating the mechanisms underlying HIPEC. There are several important strengths of the model. First, the non-invasive nature of the model allows for streamlined and reproducible application of heat. Second, the set-up of the heating device and anesthesia rig allows for higher throughput than other models that depend on individual inflow/outflow for each animal. The ability to treat and monitor multiple animals at once within the same cohort also reduces variability among animal treatments. Third, the IVIS system allows for confirmation of peritoneal carcinomatosis prior to initiation of HIPEC. In the clinical setting, HIPEC is only applied to patients with advanced EOC who have undergone an optimal resection of tumor to less than 1 cm of residual disease. In future experiments, IVIS could be used to determine the relationship between bulky abdominoperitoneal disease and HIPEC efficacy in our model. Fourth, use of mice—rather than the other mammalian species that are frequently used in animal models of HIPEC—allows for facile genetic manipulation, to determine the efficacy of HIPEC in different genetic scenarios. Additionally, the existence of immune deficient mice will allow for studies in our model that assess whether the immune system plays a role in mediating the inhibitory effects of hyperthermic cisplatin that are observed in our model of HIPEC.

While our model has several strengths, there are also several limitations. First, the use of IVIS as a way to assess tumor growth over time requires cells that express luciferase. While this is easily done in our syngeneic model using mouse cell lines, implementation in human cell lines in an immune competent mouse pose a challenge and therefore may limit the use of xenograft models in our system. Second, while the non-invasive nature of our model confers significant advantages over other animal models of HIPEC, it does differ considerably from the clinical application of HIPEC in that the mice do not undergo an open surgery. Whether surgery and the acute inflammatory response it causes contributes to the efficacy of HIPEC cannot be assessed in our non-invasive model. Third, the clinical use of HIPEC for advanced EOC involves application at the time of interval cytoreductive surgery after patients have received 3-4 cycles of platinum-based neoadjuvant chemotherapy. In our model we do not pre-treat the mice with cisplatin. Therefore, our model may not capture some important details about the effect that 3-4 cycles of platinum has on tumors and their response to HIPEC. Fourth, the current data that we show for our model does not allow for conclusions about whether HIPEC would be efficacious in the setting of platinum-sensitive vs platinum-resistant recurrent disease. Indeed, there is a great clinical need to determine effective treatments for recurrent disease, particularly platinum-resistant recurrent disease, which carries a very poor prognosis. Future studies will apply this HIPEC model on other well-established human and murine EOC to determine the impact in settings of platinum sensitive and resistant disease. This will be addressed in future experiments by pre-treating the cells with cisplatin to induce platinum resistance prior to inoculation in our model of HIPEC.

Understanding how HIPEC improves outcomes is critical to further optimize treatment for patients with advanced EOC and may shed light on targetable pathways that are currently not appreciated. HIPEC improves survival in advanced ovarian cancer patients who undergo optimal interval cytoreductive surgery [4, 28]. While the study by van Driel et al. demonstrated that HIPEC with cisplatin confers survival benefit [4], it is unclear whether this is the most effective regimen as there have not been other regimens studied in a randomized manner for this population of patients. Pre-clinical data in support of this are lacking, and our model could provide additional support for a randomized clinical trial of this regimen. Using our model, we plan to investigate this combination of HIPEC chemotherapies as well as other combinations.

In summary, we have developed a streamlined, novel, non-invasive approach to conduct HIPEC studies on mice with the ability to quantify tumor growth and assess overall animal survival. Similar to clinical outcomes with HIPEC therapy, we determined that hyperthermic cisplatin treatment of mice injected with a murine EOC cell line exhibit significant attenuation of tumor growth compared to normothermic cisplatin treatment. Studies on HIPEC animal models provide a paradigm for exploring the impact of hyperthermia and cisplatin on the tumor microenvironment and potential for application of related agents. This provides a much needed strategy to bridge the gap between *in vitro* work exploring HIPEC mechanism and the limitations of working with patient tumor within the context of standardized HIPEC therapy. The development of this noninvasive paradigm in mice provides us an opportunity to test treatment strategies for patients with advanced EOC and provide a basis for targeted drug design and development of cancer therapeutics.

## Acknowledgments

The authors would like to thank the members of the Reizes Laboratory for their insights and collaboration in completion of this work. Thanks to Dr. Yu’s Laboratory and Peng Qi from the Radiation Oncology Department in Cleveland Clinic for assistance and insights in execution of the studies.

